# Negative frequency-dependent selection maintains coexisting genotypes during fluctuating selection

**DOI:** 10.1101/729095

**Authors:** Caroline B. Turner, Sean W. Buskirk, Katrina B. Harris, Vaughn S. Cooper

## Abstract

Natural environments are rarely static; rather selection can fluctuate on time scales ranging from hours to centuries. However, it is unclear how adaptation to fluctuating environments differs from adaptation to constant environments at the genetic level. For bacteria, one key axis of environmental variation is selection for planktonic or biofilm modes of growth. We conducted an evolution experiment with *Burkholderia cenocepacia*, comparing the evolutionary dynamics of populations evolving under constant selection for either biofilm formation or planktonic growth with populations in which selection fluctuated between the two environments on a weekly basis. Populations evolved in the fluctuating environment shared many of the same genetic targets of selection as those evolved in constant biofilm selection, but were genetically distinct from the constant planktonic populations. In the fluctuating environment, mutations in the biofilm-regulating genes *wspA* and *rpfR* rose to high frequency in all replicate populations. A mutation in *wspA* first rose rapidly and nearly fixed during the initial biofilm phase but was subsequently displaced by a collection of *rpfR* mutants upon the shift to the planktonic phase. The *wspA* and *rpfR* genotypes coexisted via negative frequency-dependent selection around an equilibrium frequency that shifted between the environments. The maintenance of coexisting genotypes in the fluctuating environment was unexpected. Under temporally fluctuating environments coexistence of two genotypes is only predicted under a narrow range of conditions, but the frequency-dependent interactions we observed provide a mechanism that can increase the likelihood of coexistence in fluctuating environments.

## Introduction

Environmental variability is ubiquitous in the natural world: from daily fluctuations, to seasonal changes, to climatic variability. In contrast, evolution experiments are most commonly conducted in relatively constant environments (Lenski et al. 1991; Burke et al. 2010; Lang et al. 2013). Organisms can adapt to environmental variability in a number of ways, depending on the frequency and predictability of changes, the joint fitness landscape across environments, and genetic constraints (Beaumont et al. 2009; Kassen 2014; Botero et al. 2015). One possible outcome is selection for a single generalist genotype. In this case, selection favors genotypes with higher net fitness across the range of environmental variation. The evolution of generalists is often observed in experiments with temporally fluctuating environments (Reboud and Bell 1997; Weaver et al. 1999; Remold et al. 2008; Vasilakis et al. 2009). Generalists may be successful across environments either by acquiring mutations that are beneficial across environments, by acquiring mutations that are beneficial in one environment and have minimal cost in other environments, or by the evolution of phenotypic plasticity (Kassen 2014; Karve et al. 2016). Theory predicts that phenotypic plasticity is favored in environments where fluctuations are predictable and occur over a relatively short time span. Longer time between changes in environment instead favor repeated adaptation to the current environment (Botero et al. 2015).

Instead of a single generalist genotype, variable environments could alternatively favor the maintenance of polymorphism. In this case, multiple genotypes would be maintained in the population with each genotype having a fitness advantage in some subset of environments. Selection for multiple specialists under fluctuating environmental conditions is a possible mechanism for the maintenance of biodiversity. However, theory and experiments suggest that the conditions under which selection favors the coexistence of multiple specialists are relatively narrow. Fluctuations must be rapid enough such that neither specialist is driven extinct before the environment in which they are favored returns (Rodríguez-Verdugo et al. 2019). Furthermore, the productivity over time of the different specialists must be roughly equal so that populations of both specialists are maintained over time (Maynard Smith and Hoekstra 1980; Van Tienderen 1997).

The study of factors that promote the origin and maintenance of genetic variation despite strong selection represents an active area of research (Gloss et al. 2016; Huang et al. 2016; Edwards et al. 2018). One important question is how the period of environmental variation influences genetic polymorphism. For example, many evolution experiments are conducted by batch transfer wherein a small proportion of organisms are transferred to fresh growth media at regular intervals. Although conditions remain constant *between* growth cycles, many aspects of the environment including nutrient availability and population density can differ dramatically *during* a growth cycle. These within-day fluctuations can lead to the evolution of multiple coexisting genotypes, often via evolution of a genotype that specializes to consume metabolic by-products (Rozen and Lenski 2000; Kinnersley et al. 2014; Turner et al. 2015). Coexisting resistant and vulnerable genotypes also can emerge in predator-prey experiments where predator frequency cycles over time (Bohannan et al. 2002; Becks et al. 2010). A number of experiments have demonstrated coexistence between different pre-existing species in fluctuating environments (Legan et al. 1987; Brzezinski and Nelson 1988; Rodríguez-Verdugo et al. 2019). Outside of the evolution of cross-feeding, we are not aware of the *de novo* evolution of multiple co-existing genotypes from an experiment in which a single ancestral genotype was propagated under temporally fluctuating conditions. Indeed, even in an experiment where spatial variation in light availability selected for the evolution of coexisting genotypes, coexistence did not emerge when the environments varied temporally rather than spatially (Reboud and Bell 1997). That experiment confirmed theoretical expectations that temporal variation is less likely to promote coexistence than spatial variation (Kassen 2002).

Bacteria are useful organisms for studying the evolutionary response to fluctuating environments due to their small size, rapid reproduction, and the ability to revive frozen samples (Lenski et al. 1991). In this study, we focus on a key type of temporal variation in lifestyle for bacteria: surface-attached biofilm growth versus free-living planktonic growth. Approximately 80% of bacteria on Earth’s surface are found in biofilms (Flemming and Wuertz 2019), but planktonic growth is an important means of dispersal and allows for faster growth rates under favorable conditions. For example, during infection of the human gut, *Vibrio cholerae* transitions from a gut-attached biofilm to planktonic growth when dispersing via induced diarrhea and then back to biofilm growth upon attachment to zooplankton in water bodies (Hall-Stoodley and Stoodley 2005). Bacteria have evolved a variety of regulatory mechanisms to facilitate the switch between biofilm and planktonic modes of growth. However, growth in a constant environment may favor mutations that inactivate these regulatory mechanisms, resulting in organisms which are specialized for a particular mode of growth (Mann and Wozniak 2012; O’Rourke et al. 2015).

In this study, we report the genetic patterns of adaptation by the bacterium *Burkholderia cenocepacia* to environments that impose fluctuating selection for biofilm and planktonic modes of growth. *B. cenocepacia* is a Gram-negative bacterium typically found in agricultural soil and also an opportunistic pathogen that causes chronic pulmonary infections in individuals with the inherited disorder cystic fibrosis (Drevinek and Mahenthiralingam 2010). Replicate populations were propagated under one of three different regimes: constant biofilm, constant planktonic, or fluctuating biofilm/planktonic at weekly (~47-53 generation) intervals. We determined the identity and frequency over time of mutations in the populations via periodic whole-population, whole-genome sequencing. These data enabled us to address two main questions: 1. Does adaptation to fluctuating conditions occur via the evolution of one ecological generalist that persisted throughout the experiments, or via multiple ecological specialists that are favored during each phase of selection? 2. Does adaptation to fluctuating selection proceed by mutations in a different set of genes than those selected in constant environments?

## Methods

### Evolution experiment

We founded eight populations from a clone of *B. cenocepacia* strain HI2424, originally isolated from an onion field as an environmental isolate of the PHDC strain type that has been recovered from cystic fibrosis patients worldwide (LiPuma et al. 2002). The populations evolved for 28 days with daily transfer in M9 minimal media with galactose (GMM, 0.37 mM CaCl_2_, 8.7 mM MgSO_4_, 42.2 mM Na_2_HPO_4_, 22 mM KH_2_PO_4_, 21.7 mM NaCl, 18.7 mM NH_4_Cl, and 145 mM galactose). Populations were founded from an isolated colony grown overnight in 5 mL tryptic soy broth. Four populations were propagated under constant biofilm selection (Poltak and Cooper 2011, Figure 1A). Concurrently, four other populations evolved under a fluctuating environment regime consisting of seven days of biofilm propagation alternating with seven days of planktonic propagation (Figure 1B and 1C). During planktonic selection, 50 μL of culture was transferred to 5 mL of fresh media (6.67 generations/day). During biofilm selection, a colonized polystyrene bead was transferred to 5 mL fresh media with two sterile beads (~7.5 generations/day, Traverse et al. 2013). Before transferring, the bead was rinsed in 1 mL of phosphate buffered saline (PBS), a modification to our previously published protocol that removes residual planktonic bacteria and was predicted to strengthen selection for attachment. Following transfer to a new tube, this regime selects for bacteria that disperse from the transferred bead and attach to the new beads. Populations were incubated for 24 hours at 37 °C in a roller drum rotating at 30 rpm.

**Figure 1:**
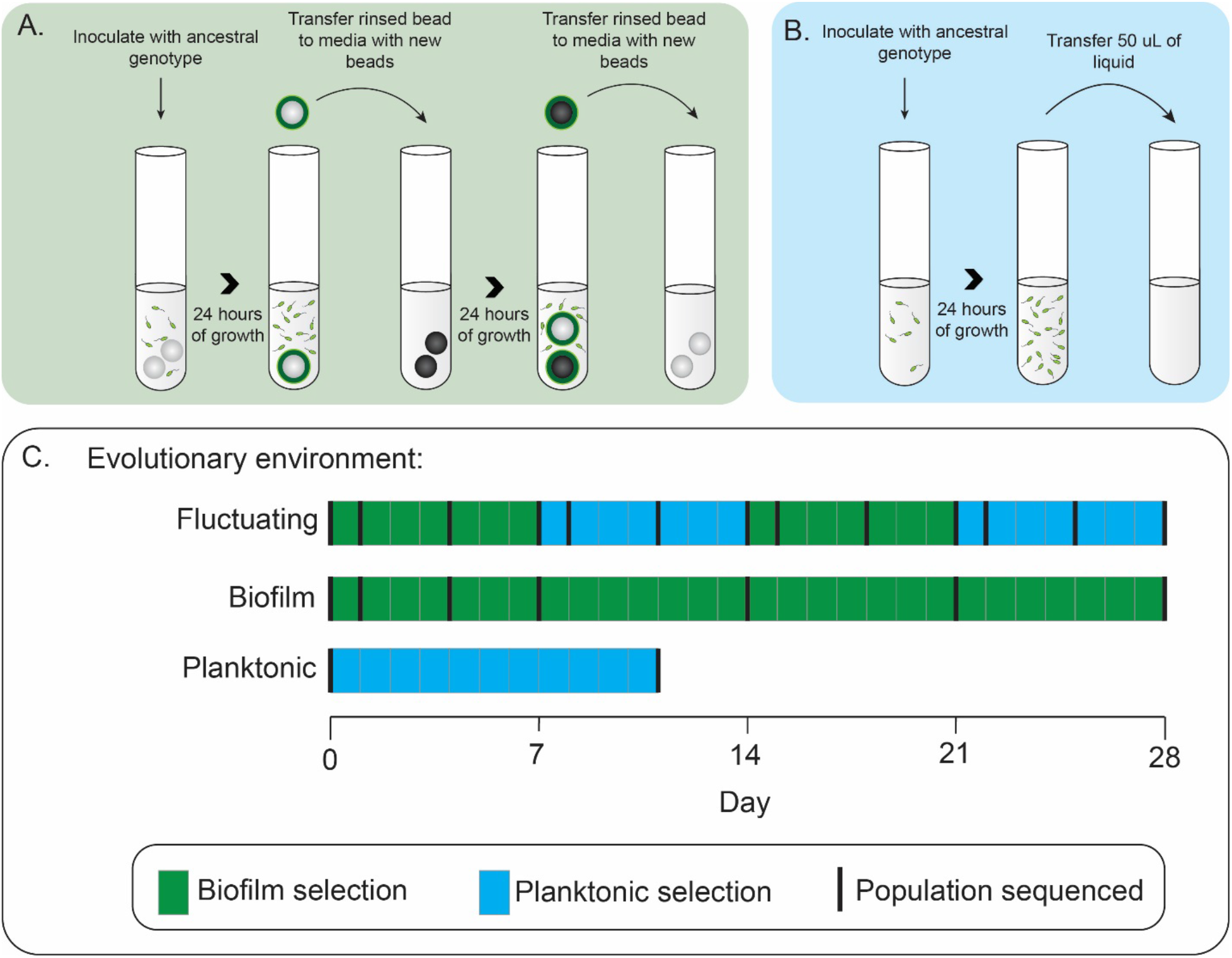
Design of evolution experiments. **A.** Under biofilm selection, a colonized bead was transferred every 24 hours (diagram modified from Turner et al. 2018). **B.** Under planktonic selection 50 μL of liquid culture was transferred every 24 hours. **C.** Populations were propagated in fluctuating biofilm and planktonic, constant biofilm, or constant planktonic environments and sequenced at indicated time points.

In the fluctuating environment, we collected population samples at days 1, 4, and 7 of each phase. In the constant biofilm environment, we collected population samples on days 1, 4, 7, 14, 21 and 28. Genomic sequencing revealed that the day 28 samples for two of these populations were contaminated so these samples were excluded from analysis. To collect biofilm population samples, one bead was rinsed in PBS, then transferred to a cryovial containing 1 mL GMM and 100 μL DMSO. To remove the bacteria from the bead, vials were vortexed before freezing at −80 °C. For planktonic samples, we transferred 50 μL of liquid culture to a cryovial containing 1 mL GMM and 100 μL DMSO and vortexed to mix.

Based on the results from the initial experiment, we additionally propagated four populations, founded by the same ancestor, under constant planktonic conditions for 11 days. We chose to evolve these populations for 11 days and compare their selected mutations with those that arose to high frequency by day 11 in the fluctuating environment. Planktonic selection was maintained in the same manner as described above for the planktonic phase of the fluctuating environment (Figure 1B). Evolution of the constant planktonic populations was performed in a different laboratory. To account for differences in trace metals in the water supply of the two labs, the following elements were added to the GMM media for these experiments: 40 μg/L Ca, 0.3 μg/L Mn, 11 μg/L K, 25 μg/L Na, 50 μg/L Zn, 0.044 μg/L Co, 0.58 μg/L Cu.

### Genome sequencing

Populations from all time points sampled were sequenced. For sequencing, populations were revived from 50 μL of frozen culture and grown under the same conditions as the evolution experiment. For biofilm samples, bacteria were removed from beads by vortexing in PBS prior to DNA extraction using the Qiagen DNeasy Blood and Tissue kit (Qiagen, Hilden, Germany). For planktonic samples, DNA was extracted using the same kit with bacteria from liquid culture. All samples were sequenced to at least 80-fold average coverage on either an Illumina HiSeq 2500 (University of New Hampshire Hubbard Center for Genome Studies) or an Illumina NextSeq 500 (University of Pittsburgh Microbial Genome Sequencing Center, University of Pittsburgh). Samples were trimmed using Trimmomatic (version 0.36, Bolger et al. 2014) and evolved mutations were identified by comparison with the ancestral *B. cenocepacia* HI2424 (GCF 000203955.1) using Breseq (version 0.28, Deatherage and Barrick 2014) with the default settings in the polymorphism mode. The threshold for detection of mutations was 0.05. We manually curated the mutations to remove false positives due to misaligned reads. We report only genes in which at least one mutation rose to 0.10 frequency or higher in at least one population sample.

### Fitness Assays

Fitness effects of evolved *wspA* F463L and *rpfR* D104G mutations were determined from clones containing representative alleles isolated from fluctuating environment populations. We focused on these two mutations because they were present at high frequency in many of our evolved populations. Whole genome sequencing confirmed the otherwise isogenic nature of the clones.

Fitness of *wspA* and *rpfR* mutants compared to the ancestor and one another was measured in both planktonic and biofilm conditions. Strains were revived from freezer stocks in 5 ml tryptic soy broth. After overnight growth, 50 μl of culture was transferred to 5 ml GMM to acclimate the strains to the competition media. After acclimation, planktonic competitions were started by inoculating 5 ml GMM with 25 ul of each competitor. Biofilm competitions were started in an identical manner, with the exception that 2 polystyrene beads were added at the time of inoculation. After 24 h, we transferred 50 ul culture to new GMM for planktonic competitions, or one bead to new GMM containing two sterile marked beads for biofilm competitions. This experimental setup closely replicated evolution conditions. Samples were collected from the competitions at day 0, 1, and 2, diluted in PBS, and plated on tryptic soy agar. For biofilm competitions, bacteria were harvested from a single bead for enumeration. To compare fitness between the *wspA* and *rpfR* mutants and the ancestor, we used a fitness-neutral, lac+ marked version of the ancestor (Poltak and Cooper 2011) and plated on tryptic soy agar supplemented with X-gal to differentiate between the lac+ and lac^−^ competitors. In competitions between the *wspA* and *rpfR* mutants, genotypes were differentiated based on their distinctive colony morphologies (small and wrinkly vs. large and smooth).

Fitness was calculated as selection rate per day:

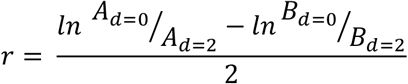

where *A* and *B* represent the densities of the two competitors and *d* indicates the timepoint. A selection rate of zero indicates that the two competitors have equal fitness. A positive selection rate indicates that competitor A is more fit, while a negative selection rate indicates that competitor B is more fit.

To measure frequency-dependent interactions between the *wspA* and *rpfR* mutants, we initiated competitions at a range of starting frequencies (approximately 0.1, 0.3, 0.5, 0.7, 0.9) by altering the volume of each competitor while maintaining total population size. We conducted these frequency-dependence competitions between the *wspA* and *rpfR* mutants in both biofilm and planktonic conditions.

In the constant planktonic environments, mutations in *rpoC* evolved in three of four populations. From one of these populations, we isolated a clone that contained only a *rpoC* mutation (T672R) and competed this isogenic mutant against a lac+-marked ancestor. In parallel, we assayed the fitness of two isogenic *rpfR* mutants (D104G and Y355D) against the lac+-marked ancestor. The *rpfR* Y355D mutation has evolved independently in multiple experiments (Traverse et al. 2013; Turner et al. 2018) and was included here as a “high fitness” *rpfR* allele. Each competition was founded with equal starting ratios of each strain and propagated planktonically.

### Growth curves and pH effects

We measured growth rate and pH tolerance to better understand the phenotypic differences between the ancestral, *rpfR* D104G and *wspA* F463L genotypes that might underlie frequency dependence between the *rpfR* and *wspA* mutants. We measured growth rate in 3% GMM in a 96-well plate on a SpectraMax plate reader (Molecular Devices, San Jose, CA). Five replicates of each strain were grown at 37° C for 24 hours, with shaking and measurement of OD_600_ every 10 minutes. Because growth in 3% GMM lowers the pH of the media to 4.1 after 24 hours, we also measured the survival of these strains in PBS at pH 4.1 and 7.0. We plated five replicate populations of each genotype at each pH on tryptic soy agar to measure the population size at 24 hours.

## Results

An evolution experiment with an environmental isolate of *B. cenocepacia* was conducted for 28 days and analyzed by population-wide, whole-genomic sequencing. Four populations were propagated under constant selection in a bead model of the biofilm life cycle (Poltak and Cooper 2011, Figure 1A), and four other populations were propagated under fluctuating selection, with one week in the biofilm model followed by one week of planktonic propagation (Figure 1B and 1C). We identified 295 mutations in these eight populations, of which 219 were nonsynonymous and 20 were synonymous base-pair substitutions. Despite the differences in selective conditions, we observed a high degree of parallel evolution at the gene level both within and between populations in both regimes (Table 1). Of 70 total genes with observed mutations, 16 had mutations at a detectable frequency in two or more biofilm-selected and fluctuating-environment populations (Tables 1 and S1). Most notably, mutations in *rpfR* (also denoted *yciR* or *pdeR* in other species), encoding a bi-functional diguanylate cyclase and phosphodiesterase as well as a sensor domain, *wspA*, encoding a transmembrane surface receptor, and *sucA*, encoding 2-oxoglutarate dehydrogenase (OGDH) were identified in all four fluctuating environment populations as well as three or more constant biofilm populations. Mutations in these genes have also been observed repeatedly in previous biofilm selection experiments with *B. cenocepacia* (Traverse et al. 2013; Turner et al. 2018). The nonsynonymous mutations in *wspA* and *rpfR* increase biofilm production by genetic de-repression, whereas mutations in *sucA* are metabolic adaptations (O’Rourke et al. 2015). Contrary to our expectations, there was no indication that the genes in which mutations were selected differed systematically between the biofilm and fluctuating environments.

**Table 1:**
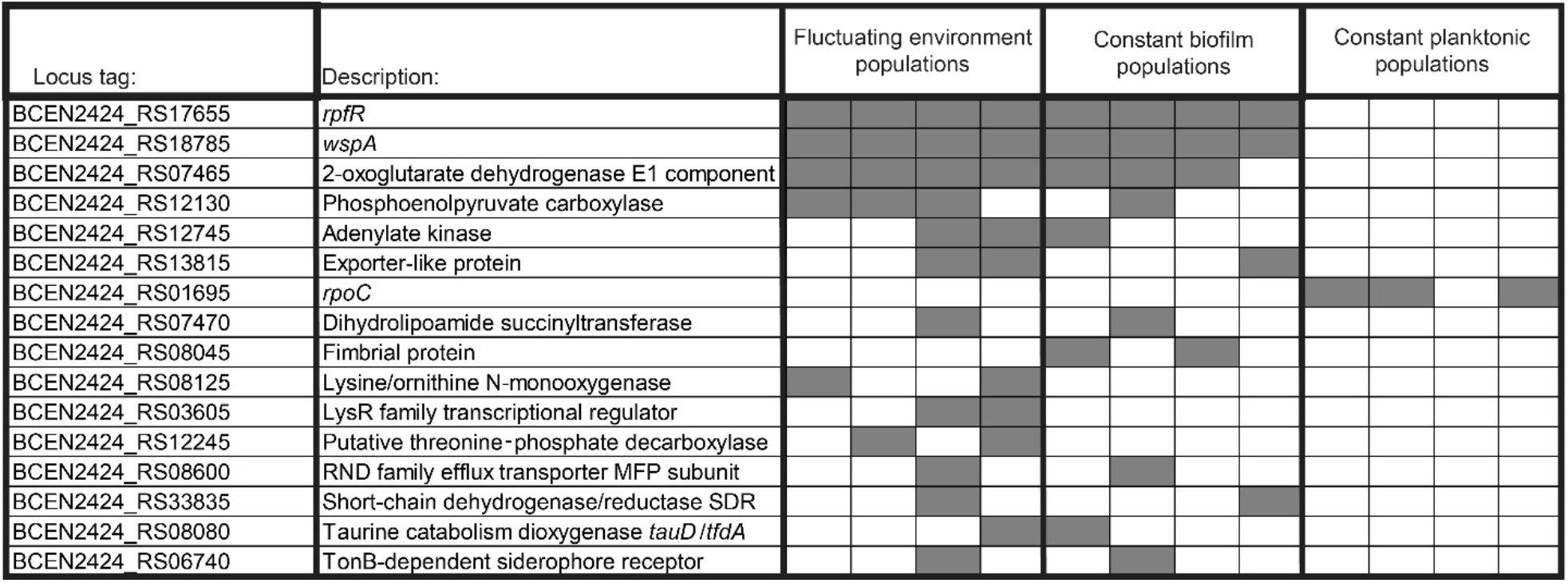
Genes in which mutations were observed in two or more populations. Each column represents one replicate population. Note that constant planktonic populations evolved for only 11 days and were sampled only at the final time point. A complete list of mutations that evolved in each population is given in Table S1.

Mutations in the biofilm-regulating genes *wspA* and *rpfR* rose to frequencies > 0.60 in all fluctuating-environment populations in at least one sample. The same mutation in *wspA* (F463L) rose to high frequency in all four constant biofilm populations and all four fluctuating environment populations (Figure 2). Given the appearance of the same exact allele and its early rise in all eight populations, we infer that the *wspA* F463L mutation was present at low frequency in the founding culture despite being undetectable by whole-population genomic sequencing. The presence of a pre-existing mutation would also help explain the rapid rise of *wspA* mutations to nearly 100% frequency in the populations. Several *rpfR* mutations were detected in multiple populations, raising the possibility of their presence at low frequency in the shared ancestral culture, though repeated independent mutations of the same *rpfR* nucleotide have been observed in prior experiments (Turner et al. 2018). Other *rpfR* alleles, however, were unique to individual populations and thus *de novo* mutations. The *rpfR* alleles ranged in identity from missense point mutations to small and large indels (Table S1).

**Figure 2:**
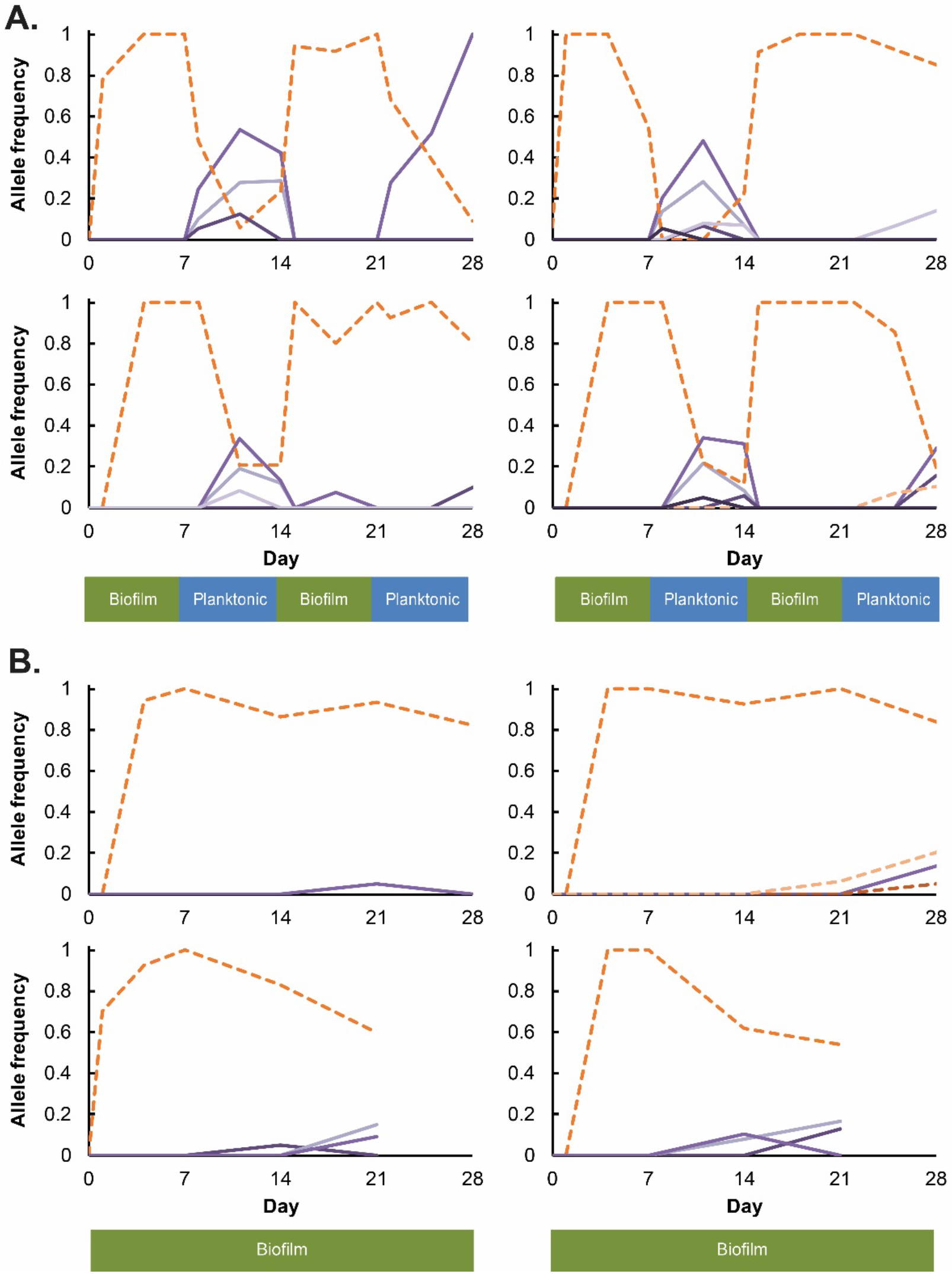
Evolutionary dynamics reveal the co-existence of *wspA* (dashed orange lines) and *rpfR* (solid purple lines) mutant genotypes in A. fluctuating and B. constant biofilm environments. Within each panel, different shades of orange and purple indicate different mutant alleles within the *wspA* and *rpfR* genes.

Remarkable parallelism also occurred in the evolutionary dynamics of the *wspA* and *rpfR* genotypes during fluctuating selection (Figure 2). The *wspA* and *rpfR* genotypes oscillated in frequency according to the environment, with *wspA* increasing during biofilm selection and *rpfR* increasing during planktonic selection. In all populations, *wspA* F463L initially spread to high frequency during the biofilm phase, but the shift to the planktonic phase selected for multiple *rpfR* genotypes that displaced the *wspA* genotype and competed with one another by clonal interference. Following the first biofilm to planktonic transition at day 7, a drastic shift in *wspA* and *rpfR* frequencies occurred in all four populations. In contrast, the frequency shifts following the second biofilm-planktonic transition were more gradual in two populations (Figure 2), possibly due to secondary mutations that were acquired on the *wspA* and *rpfR* backgrounds.

In previous evolution experiments under very similar conditions (Traverse et al. 2013; Turner et al. 2018), *rpfR* mutations were frequently selected in biofilm populations but rare in populations under planktonic selection, in contrast to the patterns seen here. We hypothesized that *rpfR* mutants had been selected during the initial biofilm phase of selection yet were outcompeted by *wspA*, and only upon transfer to the planktonic environment were *rpfR* mutants enriched because of the selective disadvantage of *wspA*. To better elucidate the selective advantages of *rpfR* mutants under planktonic selection, four replicate populations were founded from the same ancestral clone and evolved under planktonic selection for 11 days. Population sequencing failed to detect any mutations in *rpfR* above a minimum detection threshold of 5% frequency. Instead, mutations in *rpoC* (encoding RNA polymerase ß’) were detected in three of the four populations at frequencies ranging from 0.06 to 0.49. (Table 1). A clone containing only a *rpoC* mutation (T762R) was isolated from a constant planktonic population. In competitions against the ancestor, the *rpoC* mutation conferred a larger fitness benefit than both the evolved *rpfR* D104G and a high fitness *rpfR* mutation that repeatedly evolved in previous experiments (Y355D, Fig. 3). This result suggests that though *rpfR* is beneficial in planktonic conditions, its fitness effect is less than that of other available mutations – such as *rpoC* – and thus it fails to rise to a detectable frequency during constant planktonic selection.

**Figure 3:**
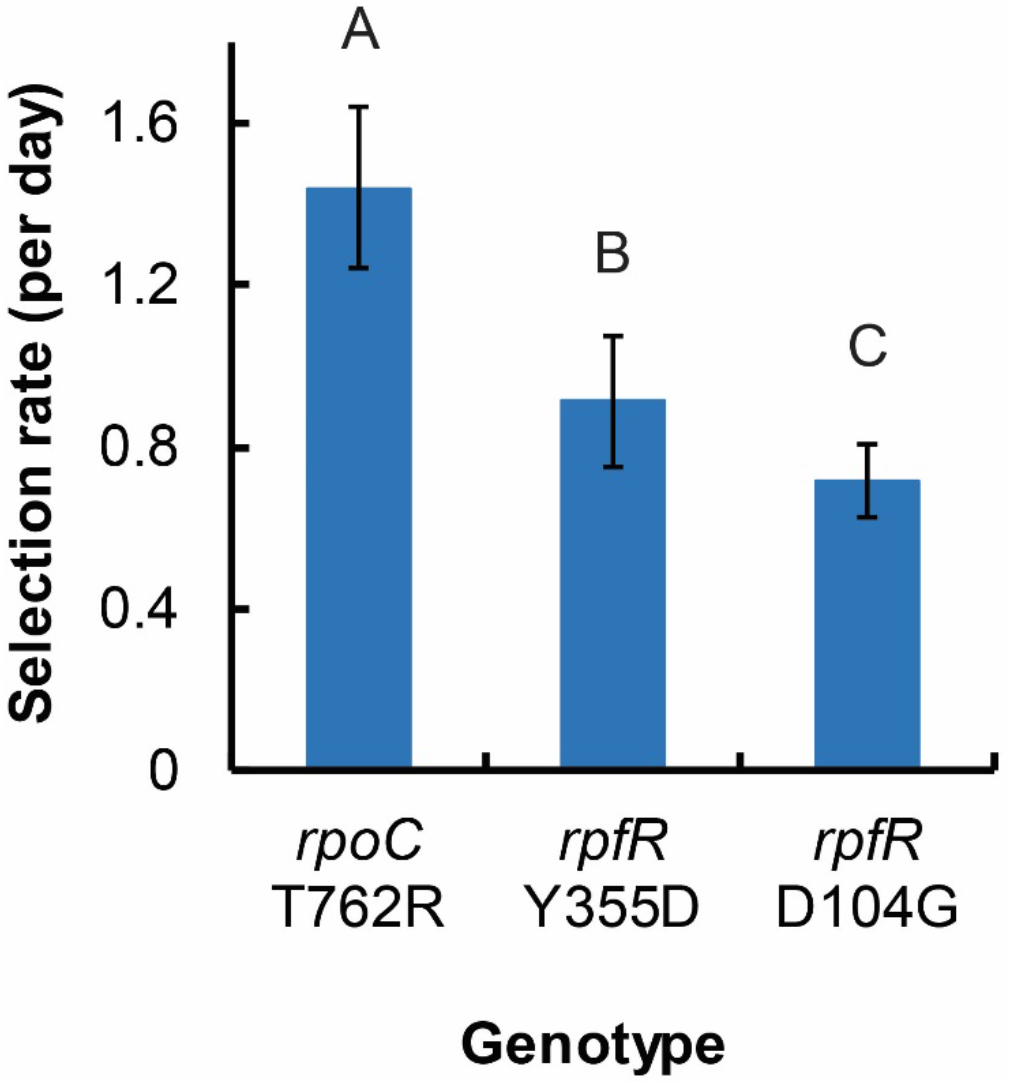
The planktonic fitness advantage of the *rpfR* mutants is less than that of the *rpoC* T762R mutant isolated from a constant planktonic population (mean ± 95% confidence interval, ANOVA with Tukey’s post-hoc test F=47.8, *p* < 10^−5^, bars with different letters are significantly different from each other).

Next, we explored the ecological basis of the observed coexistence between the *wspA* and *rpfR* genotypes. Clones containing only the *wspA* (F463L) or the *rpfR* (D104G) mutation were isolated from evolved fluctuating populations. From the observed evolutionary dynamics of the fluctuating environment populations (Fig. 2A), we expected the *rpfR* mutant to have higher fitness in the planktonic environment and the *wspA* mutant to have higher fitness in biofilms. Indeed, the ancestor was outcompeted by the *rpfR* mutant in planktonic conditions and by the *wspA* mutant in biofilm conditions (Fig. 4A). The *wspA* mutant exhibited a fitness tradeoff in planktonic conditions, consistent with the observed evolutionary dynamics. Surprisingly, however, the *rpfR* mutant exhibited a significant fitness advantage over the ancestor in biofilm conditions, to a similar extent as the *wspA* mutant. Further, when competed head-to-head in biofilm conditions, the fitness of the *rpfR* mutant was indistinguishable from the *wspA* mutant. These data suggest that the *rpfR* mutation provides a fitness advantage in both environments, prompting the question of why *wspA* genotypes dominated the *rpfR* genotypes during the biofilm phase of the fluctuating environment regime.

**Figure 4:**
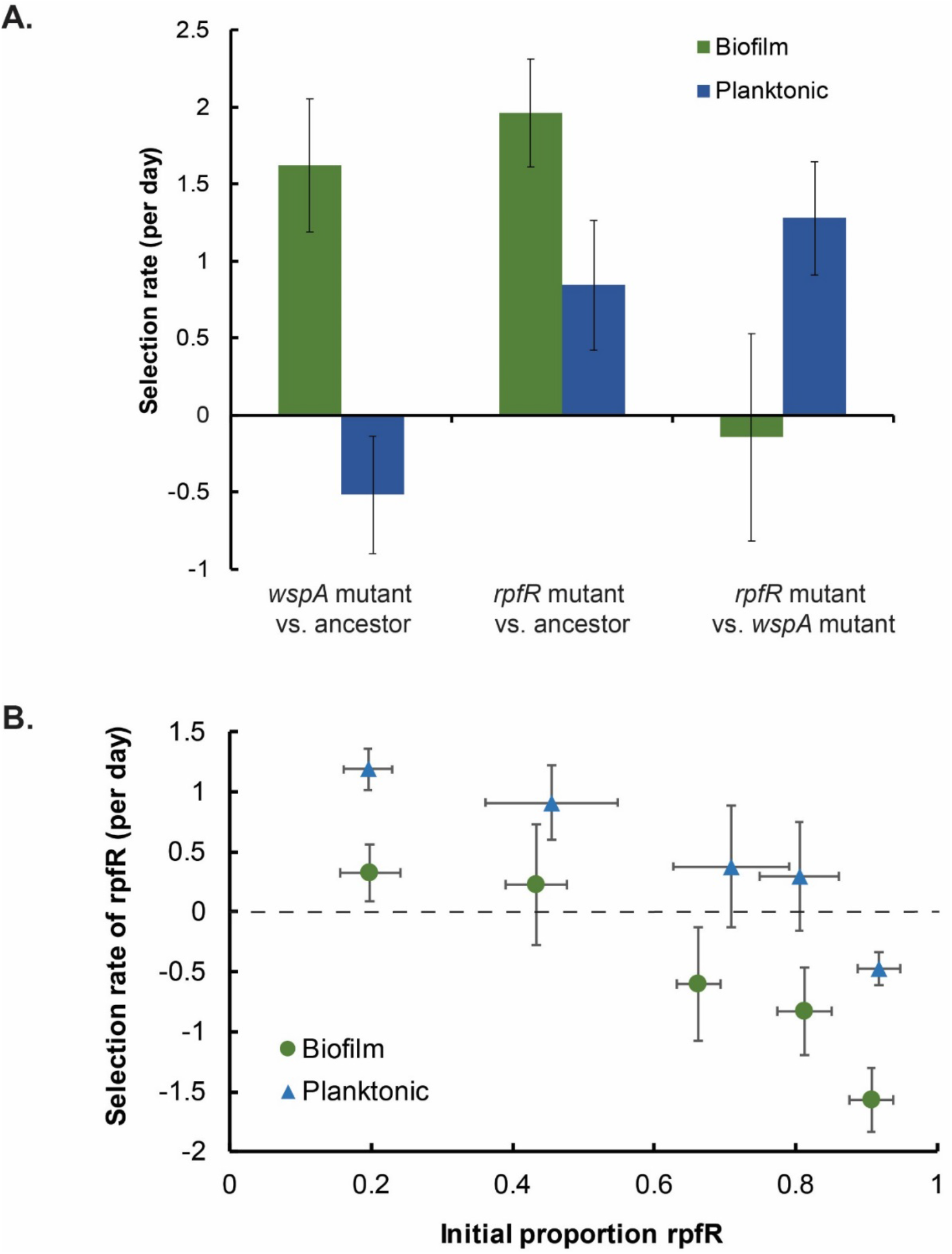
The *rpfR* (D104G) and *wspA* (F463L) mutants exhibit negative frequency-dependence under both biofilm (green) and planktonic (blue) conditions. A. Fitness (mean ± 95% confidence interval) of *wspA* F463L and *rpfR* D104G genotypes in pairwise competitions with equal starting ratios. B. Fitness at a range of starting ratios.

A potential explanation for mutant coexistence could involve frequency-dependent selection. Whereas the original fitness assays of evolved mutants were performed by mixing strains in equal ratios, we investigated whether starting ratios influenced fitness by combining mutants across a range of frequencies. Negative frequency dependence – or advantage-when-rare -- is evident in competitions between the *wspA* and *rpfR* mutants in both planktonic and biofilm conditions (Fig. 4B). However, the equilibrium frequency – the frequency at which both strains are equally fit – shifted between environments, with *rpfR* having a higher equilibrium frequency under planktonic conditions and a lower equilibrium frequency under biofilm conditions.

A possible explanation for negative frequency dependence is ecological differentiation between the strains. Under planktonic conditions, the *rpfR* D104G mutant grew more quickly and to a higher final density than the ancestral genotype (Fig. S1A). In contrast, the *wspA* F463L mutant grew more slowly and to a lower final density than either the ancestor or the *rpfR* mutant. However, the *wspA* mutant exhibited increased survival at pH 4.1, which is the pH of GMM following 24 hours of growth and acidification of the environment through metabolic by-products (Fig. S1B), whereas there was no difference in survival at pH 7.0. This result suggests that broader niche breadth of the competitor with inferior growth rate may maintain genotype coexistence.

## Discussion

Here we report the genetic basis of adaptation and evolutionary dynamics of replicate populations of *B. cenocepacia* under fluctuating selection for biofilm and planktonic growth. Rather than favoring a single genotype with the ability to succeed across both biofilm and planktonic conditions, the fluctuating environment selected for two co-existing lineages, each of which had an advantage during a particular phase of the experiment. In fluctuating populations, we observed repeated shifts in frequency between two lineages (Fig. 2), each with a mutation in a different biofilm-regulatory gene (*wspA* and *rpfR*). When competed against each other, *wspA* and *rpfR* isogenic mutants exhibited stable negative frequency-dependence, such that both mutants were able to coexist in each environment (Fig. 4B). However, the frequency of the stable equilibrium shifted depending on the environment. Under biofilm conditions *wspA* genotypes dominated, forcing *rpfR* mutants below detection, whereas under planktonic selection conditions the *rpfR* genotypes reached a higher frequency owing to the fitness cost of *wspA* in this condition.

Frequency-dependent coexistence between the two mutants in planktonic conditions may be driven in part by the acidification of the media during growth to a pH of 4.1. Despite an initial growth rate advantage of the *rpfR* mutant, the *wspA* mutants exhibit greater survival in acidic conditions caused by metabolic byproducts of growth in the galactose minimal medium (Fig. S1). The greater biofilm production of *wsp* pathway mutants relative to *rpfR* mutants (Poltak and Cooper 2011; Traverse et al. 2013) may enable greater tolerance to the stress of low pH. Under biofilm conditions, *wspA* and *rpfR* mutants have been shown to form distinct biofilm structures, with the *wspA* genotype attaching early, tightly, and directly to the plastic bead during the biofilm phase while *rpfR* mutants tend to attach later and adhere to both the plastic bead and to other adherent cells (Poltak and Cooper 2011; Ellis et al. 2015). These distinctions provide physiological explanations for their frequency-dependent interactions in the biofilm environment, which requires both biofilm growth and planktonic dispersal during each cycle.

Stable negative-frequency dependent coexistence with shifting proportions of *wspA* and *rpfR* can broadly explain the dynamics observed in the evolution experiment, where *wspA* increased in frequency during biofilm selection, while *rpfR* increased in frequency during planktonic selection. However, the frequencies observed in the evolution experiment differed from those predicted by the competitions between individual strains. Specifically, *rpfR* mutations were undetectable during biofilm selection whereas the competition experiments suggested that *rpfR* should have an equilibrium frequency of more than 20% in biofilm conditions. These differences could arise due to the “head start” of *wspA* mutations in this experiment and later be influenced by additional adaptive mutations arising in *wspA* and *rpfR* genotypes that improved lineage fitness. The evolutionary head start of the *wspA* mutations is partly due to biofilm selection occurring first in our fluctuating environments. In addition, it appears likely that the *wspA* mutation observed in our populations was present at a low frequency in the founder of our experimental populations. This too could have contributed to an evolutionary head start for the *wspA* lineage.

Our results suggest that coexistence between *rpfR* and *wspA* mutants is ecologically stable, with the two genotypes coexisting under both biofilm and planktonic conditions. However, it is unclear if coexistence would be stable over longer evolutionary time frames. In most populations, *wspA* genotypes reached a higher frequency during the second week of planktonic selection than the first, leading us to speculate whether a single genotype would eventually fix given enough time. The increased frequency of *wspA* lineages may be explained by secondary mutations that compensated fitness under planktonic conditions. Furthermore, epistatic effects of the initial beneficial mutations could increase or decrease access to subsequent beneficial mutations, producing a scenario in which *wspA* mutants adapt more rapidly to planktonic conditions than *rpfR* mutants adapt to biofilm conditions. Ultimately, however, the success of the *wspA* genotypes may simply result from its presence in the standing variation of the ancestral culture and its dominance in the early phase of the fluctuating regime, and hence its earlier access to secondary, beneficial mutations.

Surprisingly, although the frequencies of mutations within populations differed, there was no clear difference between fluctuating and constant biofilm environments in the identity of the genes in which mutations were selected. This could be simply explained by the requirement that bacteria disperse and attach to the new plastic bead in our biofilm model, essentially producing daily periodicity between biofilm and planktonic growth. Somewhat related, an inconsistency in the strength of selection imposed during each phase of the fluctuating environment may partially explain why the mutational spectra of the constant biofilm and fluctuating environment populations appear indistinguishable. Assuming stringent biofilm selection, a lineage must acquire a mutation that improves fitness in the biofilm phase in order to reach the subsequent planktonic phase, at which point available mutations that also afford an advantage in planktonic condition would begin to dominate. This appears to be the case for the *rpfR* mutations that enable their lineages to survive the stringent biofilm selection and then thrive upon the shift to the milder planktonic selection.

In contrast, no mutations were observed in common between the fluctuating populations and the 11-day constant planktonic populations. No mutations in *rpfR* were observed in any of the planktonic populations, even though *rpfR* mutations rose to high frequencies by day 11 in all four fluctuating environment populations. Although *rpfR* mutants are more fit than the ancestor in planktonic conditions (Fig. 4), they are less fit than the *rpoC* mutations that were observed in the planktonic-selected populations (Fig. 3). There are multiple examples of mutations in subunits of RNA polymerase selected during bacterial evolution experiments selecting for rapid growth in minimal media, which provides evidence of their specific advantages in planktonic, serially-diluted culture conditions (Barrick et al. 2010; Conrad et al. 2010; Rodríguez-Verdugo et al. 2014). *rpfR* mutations were also rarely observed under planktonic selection in previous similar evolution experiments (Traverse et al. 2013; Turner et al. 2018). These results further support the conclusion that the prevalence of *rpfR* mutations in the planktonic phase of the fluctuating environment was driven by their combined fitness across both biofilm and planktonic environments, rather than solely by their advantage in planktonic environments.

Adaptation to variable environments can occur through evolution of generalists, phenotypic plasticity, or coexistence of specialist genotypes. Here, the evolution of *B. cenocepacia* under fluctuating planktonic or biofilm forms of growth selected for two coexisting genotypes, each having an advantage in a different phase of the experiment. In contrast, previous evolution experiments involving temporally fluctuating selection selected a single lineage of generalist mutants (Reboud and Bell 1997). Furthermore, theory indicates that coexistence of genotypes in temporally fluctuating environments is less likely because it requires equal productivity of genotypes over time (Maynard Smith and Hoekstra 1980; Van Tienderen 1997). In the current experiment, coexistence was facilitated by negative frequency-dependent interactions in which both genotypes were able to stably coexist (at least over ecological timescales) in both environments. The environmental shifts simply altered the expected frequencies of each genotype. Our results raise the possibility that frequency-dependent interactions could promote the likelihood and stability of coexisting genotypes as an outcome of adaptation to non-constant environments. Recent work in microbial experimental evolution has suggested that stable frequency-dependent coexistence may be a more common than previously expected (Good et al. 2017), and population-genetic surveys of bacteria colonizing humans also indicate that frequency-dependent dynamics may be common (Silva et al. 2016; Zhao et al. 2019). This growing evidence suggests that it would be valuable to develop theory considering the effects of frequency-dependent coexistence on adaptation to variable environments, as well as greater investment in studies of standing genetic diversity in populations that could be maintained by environmental periodicity (Corander et al. 2017).

## Acknowledgements

We thank D. Snyder, N. Phillips, N. Rouillard and K. Koerner for assistance in the laboratory. This work was funded by NIH (R01GM110444) and NASA Astrobiology Institute (NAI CAN-7 NNA15BB04A) grants to VSC.

## Data accessibility statement

All data and R scripts will be made available on Dryad. Raw sequencing reads will be submitted to NCBI SRA.

## Author contributions

SWB and VSC designed the research. SWB conducted the original experiment and CBT conducted subsequent laboratory research. CBT and VSC wrote the manuscript with input from all authors.

---

**Figure S1:**
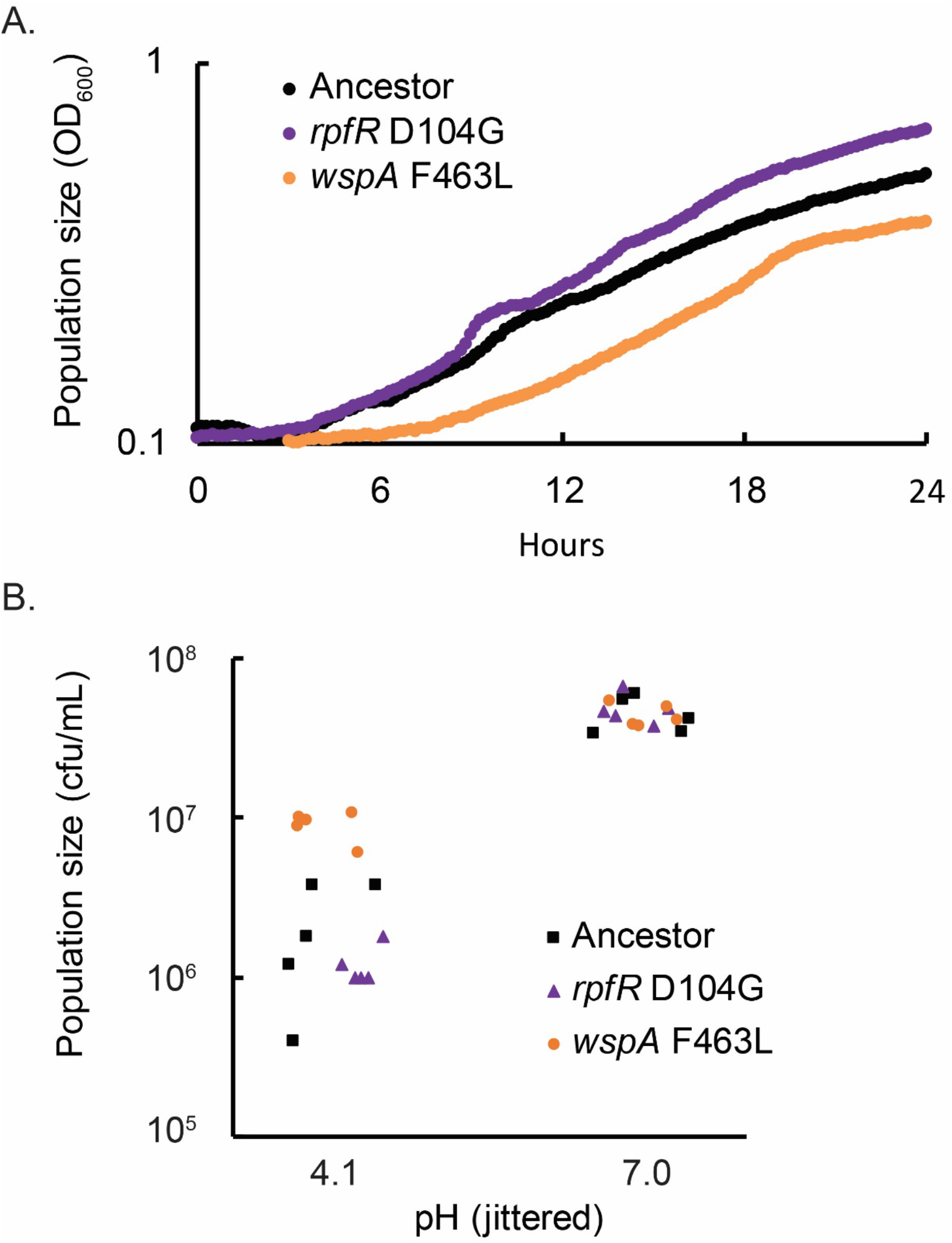
Growth and survival differences between *rpfR* and *wspA* mutants. A. The *rpfR* D104G mutant grew more quickly than the *wspA* F463L mutant in evolution media. B. The *wspA* F463L mutant had higher survival in pH 4.1 PBS for 24 hours than the *rpfR* D104G mutant (ANOVA with Tukey’s post-hoc test F=46.2, *p* < 10^−5^), while there were no differences between genotypes in survival at pH 7 (ANOVA F=0.3, *p* = 0.77).

